# Reorganizations of latency structure within white matter from wakefulness to sleep

**DOI:** 10.1101/2021.08.25.457605

**Authors:** Bin Guo, Fugen Zhou, Guangyuan Zou, Jun Jiang, Qihong Zou, Jiahong Gao

## Abstract

Previous studies based on resting-state fMRI (rsfMRI) data have revealed the existence of highly reproducible latency structure, reflecting the propagation of BOLD fMRI signals, in white matter (WM). Here, based on simultaneous electroencephalography (EEG) and functional magnetic resonance imaging (fMRI) data collected from 35 healthy subjects who were instructed to sleep, we explored the alterations of propagations in WM across wakefulness and nonrapid eye movement (NREM) sleep stages. Lagged cross-covariance was computed among voxel-wise time series, followed by parabolic interpolation to determine the actual latency value in-between. In WM, regions including cerebellar peduncle, internal capsule, posterior thalamic radiation, genu of corpus callosum, and corona radiata, were found to change their temporal roles drastically, as revealed by applying linear mixed-effect model on voxel-wise latency projections across wakefulness and NREM sleep stages. Using these regions as seeds, further seed-based latency analysis revealed that variations of latency projections across different stages were underlain by inconsistent temporal shifts between each seed and the remaining part of WM. Finally, latency analysis on resting-state networks (RSNs), obtained by applying k-means clustering technique on group-level functional connectivity matrix, identified a path of signal propagations similar to previous findings in EEG during wakefulness, which propagated mainly from the brainstem upward to internal capsule and further to corona radiata. This path showed inter-RSN temporal reorganizations depending on the paired stages between which the brain transitioned, e.g., it changed, between internal capsule and corona radiata, from mainly unidirectional to clearly reciprocal when the brain transitioned from wakefulness to N3 stage. These findings suggested the functional role of BOLD signals in white matter as a slow process, dynamically modulated across wakefulness and NREM sleep stages, and involving in maintaining different levels of consciousness and cognitive processes.

## Introduction

Recent studies utilized latency analysis to investigate spatiotemporal organizations of the brain. Results from various signal modalities, including EEG (Nir et al. 2011; Mitra et al. 2016), MEG (De Pasquale et al. 2010) and BOLD fMRI (Marek et al. 2018; Mitra et al. 2014; Raut et al. 2021), indicated that coordination of brain areas occurred on the time scales spanning from milliseconds to tens of seconds by various methods of latency analysis. While conventional zero-lag temporal correlation (referred to as functional connectivity, FC) analysis focused on detections of concurrent spontaneous fluctuations, assumed to persist over minutes (Biswal et al. 1995; Fox et al. 2007), latency analysis complemented the FC analysis in that it advanced our understanding of the spatial-temporal organizations of spatially segregated brain areas, e.g., resting-state networks (RSNs).

In human gray matter (GM), comprehensive latency analysis has been performed to investigate the lead/lag relationships within cerebral cortex (Mitra et al. 2014), between thalamus and cerebral cortex (Mitra et al. 2015), between hippocampus and cerebral cortex (Mitra et al. 2016), and within cerebellum (Marek et al. 2018), suggesting that latency structure of spontaneous fluctuations in GM BOLD fMRI signals manifested as a slow neural progress varying across physiological states. On the contrary, BOLD signals from white matter (WM) have long been regarded as artefacts, leading to the ignorance of the functional role of WM BOLD signals. However, a recent complemented study (Guo et al. 2021a) on a large cohort of youngsters revealed the existence of highly reproducible latency structure of BOLD fMRI signals in WM. Variations of the latency structure were found to be associated with different sensory states (eye closed vs eye open). Also, growing evidence converged to support the conclusion that fluctuations in WM BOLD signals also encoded neural activity (Gawryluk et al. 2014; Ding et al. 2018; Grajauskas et al. 2019; Gore et al. 2019). Providing further evidence, a recent study using joint MRI/PET dataset (Guo et al. 2021b) found correlations between FC and fluorodeoxyglucose and also between FC and the fractional amplitude of low frequency fluctuations in WM bundles, indicating that BOLD signals in WM reflected variations in metabolic demand associated with neural activity.

With the observations above, we hypothesized that the propagations of BOLD fMRI signals in WM also behaved as a slow process involving in sustaining different levels of consciousness and cognitive processes by dynamically modulating directions of signal propagations among regions across wakefulness and nonrapid eye movement (NREM) sleep stages. To test this hypothesis, lagged cross-covariance (Mitra et al. 2014; Raut et al. 2020; Guo et al. 2021a) and FC analysis were both performed on BOLD fMRI signals from a simultaneous EEG-fMRI dataset. Latency analyses were performed in three different aspects. First, latency projections (Mitra et al. 2014; Guo et al. 2021a) showed significant reorganizations of latency topographies, which suggested reorganized apparent propagations of BOLD signals in WM regions across wakefulness and NREM sleep stages in an average sense. Second, using the regions found in the first step as seeds, we further calculated seed-based latency maps, reflecting latency value of each voxel with respect to the reference seed, and found that the reorganizations of latency topographies were driven by inconsistent shifts of latencies shared by different regions in WM across wakefulness and NREM sleep stages. Third, latency analysis was performed on WM resting-state networks (RSNs), derived by applying K-means clustering technique on voxel-wise FC matrix. A path of signal propagations, starting from brainstem upward to internal capsule (IC) and further upward to corona radiata (CR), was identified during wakefulness at this level of latency analysis. In descent to deep sleep, the path changed to exhibit different patterns of inter-RSN temporal organizations depending on the paired stages between which the brain transitioned. These findings validated our assumption to regard the propagations of BOLD fMRI signals in WM as a slow process, dynamically modulated across wakefulness and NREM sleep stages, involving in maintaining different levels of consciousness and cognitive processes.

## Materials and Methods

### Subjects

Resting-state data from 35 right-handed healthy subjects (16 males, mean age 21.7±3.2 years, age range:18-29 years) were recruited to participate in the experiment. None of the subjects had a history of brain injury, psychiatric or neurological disease, psychoactive drug use and drug or alcohol abuse. During experiment days, all subjects were prohibited from consuming alcohol or caffeine. Informed consent was provided by all subjects prior to the experiments.

### Data acquisition

Simultaneous EEG-fMRI data were acquired using a 3T Siemens Prisma MRI scanner (Siemens Healthineers, Erlangen, Germany) with a 64-channel MR-compatible EEG system (Brain Products, Munich, Germany), and the sampling rate was 5000 Hz. During the experiment, the subjects were required to lie quietly and their heads were retrained by cushions. The recording montage consisted of 64 channels, including 57 EEG channels, two reference channels (A1, A2), two electromyography (EMG) channels, two electrooculography (EOG) channels and one electrocardiogram (ECG) channel, and the channels were arranged according to the international 10/20 system. The resistance of the reference and ground channels was maintained at less than 10 kΩ, and the resistance of the other channels was maintained at less than 20 kΩ. The resistance of all the channels was confirmed to meet the criteria before the MR scanning session started and after it ended.

Simultaneous EEG-fMRI data and high-resolution anatomical images were acquired using a gradient echo planar imaging (EPI) sequence (repetition time (TR) = 2000 ms, echo time (TE) = 30 ms, flip angle (FA) = 90°, number of slices = 33, slice thickness = 3.5 mm, gap = 0.7 mm, matrix = 64 × 64, and inplane resolution = 3.5 × 3.5 mm2) and a three-dimensional magnetization-prepared rapid gradient echo T1 weighted sequence (TR = 2530 ms, TE = 2.98 ms, inversion time = 1100 ms, FA = 7°, number of slices = 192, matrix =512 × 448, and voxel size = 0.5 × 0.5 × 1.0 mm3), respectively. During EEG-fMRI scanning, the subjects were asked to close their eyes and try to sleep. Scanning ended when the largest number of volumes (4096 volumes for the EPI sequence in our scanner) was recorded or the subjects were completely awake after sleeping and were unable to fall asleep again.

### EEG preprocessing and sleep stage scoring

EEG data were preprocessed using BrainVision Analyzer 2.1 (Brain Products, Munich, Germany). MR gradient artifacts were corrected utilizing the average artifact subtraction method. For the ballistocardiogram artifacts, the software-misidentified R peaks were manually adjusted after semiautomatic R peak detection. The average artifacts were subtracted from the EEG data after transferring ECG R peaks to the EEG data over a selectable time delay. Next, the data were downsampled to 500 Hz, referenced to the mean values of reference channels (A1 and A2). Last, the data were temporally filtered (10–100 Hz for electromyogram channels and 0.3–35 Hz for the other channels) (Zou et al. 2021). Sleep stage scoring was performed visually by two experienced technicians for every 30-s duration of preprocessed EEG data based on the American Academy of Sleep Medicine criteria, and the scoring result of one technician was double checked by the other. Sleep EEG recordings were divided into five stages, including eyes-closed wakefulness (Wake), N1, N2, N3 and rapid eye movement (REM) sleep.

### Functional and anatomical preprocessing

The fMRI data were grouped into epochs of 5 minutes based on the EEG sleep stage scoring. Each session corresponding to wake state or a specific sleep stage. Epochs shorter than 5 minutes were excluded. Also, subjects with large head motion (>3 mm in translation or >3° in rotation) were excluded. Under these criteria, the fMRI data of the 35 subjects were grouped into 40 sessions in wakefulness (12 subjects), 26 sessions in N1 (17 subjects), 104 sessions in N2 (24 subjects), and 104 sessions in N3 (24 subjects). No 5-minute continuous REM sleep stages were identified since NREMs occupied the majority of sleep during the first half of the night.

Images were preprocessed using the statistical parametric mapping software package SPM12 (www.fil.ion.ucl.ac.uk/spm/software). First, the images were corrected for slice timing and head motion. Second, T1-weighted images were segmented into gray matter, white matter, and cerebrospinal fluid using the New Segment utility embedded in SPM12, and all these images were registered to the mean BOLD image output by the motion correction procedure. Third, the BOLD images were normalized into the Montreal Neurological Institute (MNI) space (3mm), along with the coregistered T1-weighted images as well as the gray matter and white matter segments. Fourth, linear trends from the BOLD images were removed to correct for signal drifting. Fifth, mean signals from the cerebrospinal fluid mask and whole-brain mask were regressed out as nuisance covariates. After the nuisance regression, the temporal signals were low-pass filtered to retain frequencies between 0.01∼0.1Hz. Finally, BOLD images were spatially smoothed with FWHM = 6 mm isotropic Gaussian filter, where smoothing was performed separately on the white matter and the gray matter of each subject, to avoid mixing in-between signals.

### Creation of group-level white matter mask

Following the above preprocessing steps, the white matter mask of each subject was normalized to the MNI space and resampled to isotropic resolution of 3 mm. To obtain the masks for group-wise analysis, we used the segments for each subject. The subject-level white matter mask was generated using a threshold value of 0.8, and the strategy of majority voting was applied to generate group-level mask. Voxels with a voting rate above 90% will be included in the group-level mask, resulting in a group-level mask consisting of deep white matter voxels, which contained 11523 voxels.

### Estimation of time delay between BOLD time series

Given time series of two voxels, *x*_1_(*t*) and *x*_2_(*t*), a lagged cross-covariance was computed as,

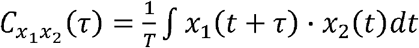

where *τ* is the latency (in units of time), and T is the integration interval, a constant in our experiment. The value of *τ* at which 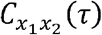 exhibiting an extremum defines the temporal latency (equivalently, delay) between signals *x*_1_ and *x*_2_ (Konig 1994). To determine the exact latency time value at the resolution finer than the scale of time of repetition, a parabolic interpolation approach was used, as implemented previously (Mitra et al., 2014; Laumann et al., 2017; Raut et al., 2020).

### Generation of latency matrices and latency projections

Latency matrix, with each entry representing the temporal latency between paired voxels, necessarily anti-symmetric, was computed for each subject and then averaged across each stage to obtain a stage-specific latency matrix of size 11523×11523. And the latency projection is obtained by averaging over the rows of the latency matrix (Nikolic 2007). The latency projection represents the relative timing of each voxel with respect to the remaining voxels selected. Latency projections were derived from each stage-specific latency matrix.

### Generation of seed-based latency maps

BOLD time series of voxels from a selected seed (sub-region of a mask) was averaged and regarded as a reference for remaining voxels to estimate in-between latencies, resulting in a one-dimensional seed-based latency map.

### Generation of RSNs in white matter

Previous studies on GM demonstrated that the topography of RSNs, especially the default mode network, was generally preserved across sleep stages (Horovitz et al. 2009; Larson-Prior et al. 2009; Tagliazucchi et al. 2013b). Preconditioning on this observation (see Discussion), we applied traditional clustering technique, K-means (Peer et al. 2017; Guo et al. 2021), on group-level voxel-wise FC matrix, obtained from the N2 group of subjects since more subjects were included in this group, to identify the RSNs in white matter.

### Latency matrices of inter- and intra-RSNs

Similar operations (Guo et al. 2021; Mitra et al. 2015) were performed to derive inter- and intra-RSN latency matrices. Briefly, the group-level latency matrix was reordered by network membership to group voxels from the same RSN in the same latency block. For each latency block, rows and columns were rearranged according to the temporal orders within corresponding networks.

### Statistical analysis of latency projections

Linear mixed-effect analysis (Chen et al., 2013) was adopted to study the main effect of different sleep stages, along with the paired effects in between. The modeling program, 3dLMEr from AFNI (Cox 1996), was utilized to build the statistical model, in which variates of sleep stages (Wake, N1, N2, and N3), age, gender, and years of education were included. Cluster correction was employed for evaluating both main and paired effects. Threshold of cluster size of statistical significance was determined by autocorrelation function modeling combined with a voxel-level threshold of p < 0.001. To be specific, we applied 3dFWHMx on preprocessed fMRI data to estimate the parameters for autocorrelation function (ACF), followed by 3dClustSim with 10000 iterations for identification of significant cluster-size, within which a corrected significance level was thresholded at p < 0.01 and uncorrected voxel-level at p < 0.001 (Eklund et al. 2016).

## Results

### Main and paired effects of stages on latency projections

#### Main Effect

The latency projections of different stages were shown in Figure 1-A. Main effect of stages on the distribution of latency projection was shown in Figure 1-B. Regions showing statistical significance included cerebellar peduncle (CP), IC, posterior thalamic radiation (PTR), genu of corpus callosum (GCC), and CR.

**Figure 1.**
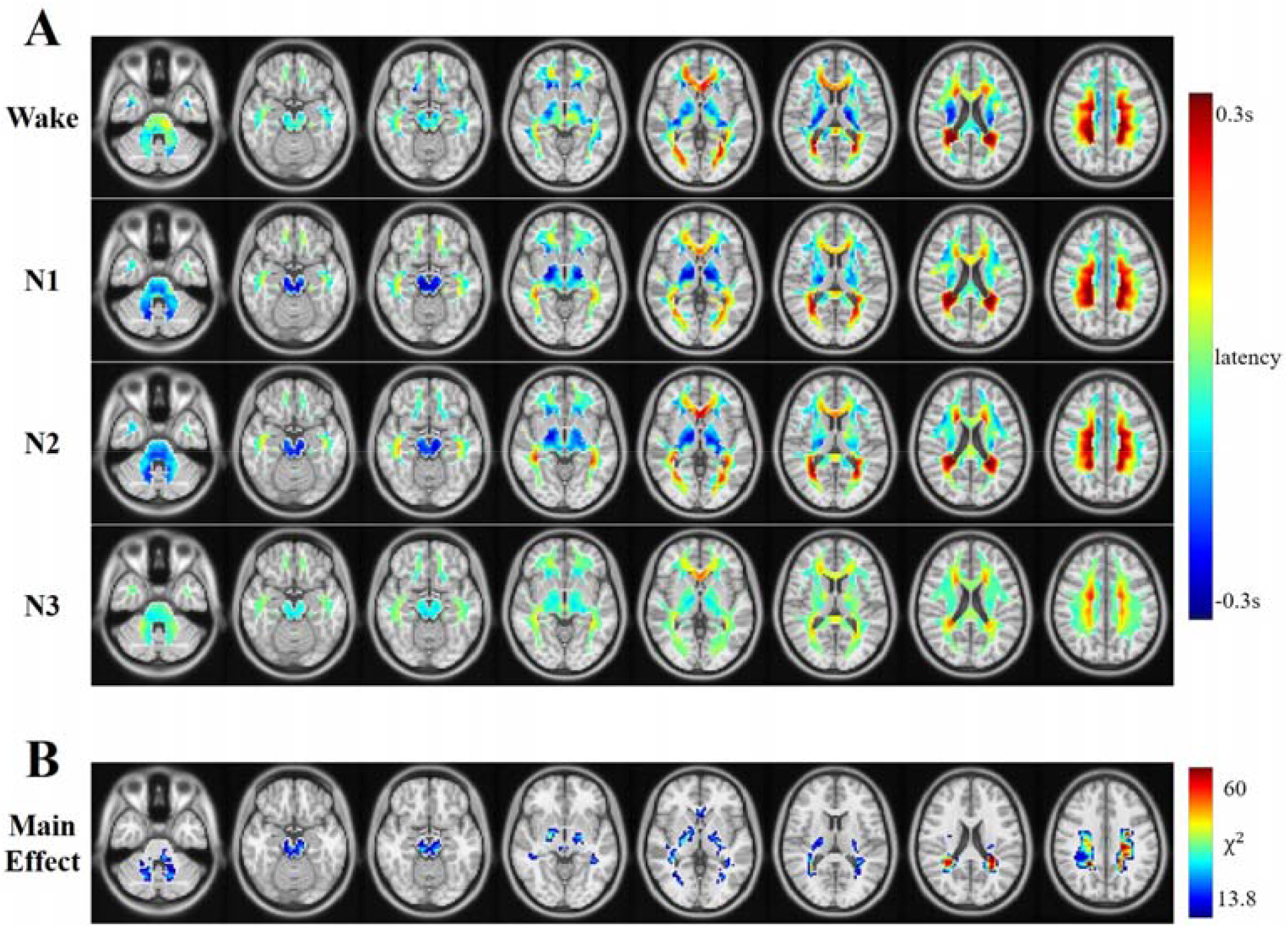
(A) Latency projections in white matter across wakefulness and NREM sleep stages. Same colormap is applied to all the latency projections. (B) Distributions of main effect of different stages on latency projection, wherein >13.8 and p<0.01 with cluster-wise correction.

#### Significant Clusters of the paired effects

Post-hoc t-test was performed on each of the paired stages to explore the actual paired stage that drove the variations of latency projections. Only voxels contained within the main effect regions were included for statistical analysis. In post-hoc t-test, significant regions were defined by thresholding at p < 0.001, wherein clusters of size larger than 20 (corresponding to p < 0.01, corrected) were reported and kept for further seed-based latency analysis.

#### Paired effect of Wake and N1

Figure 2-A displays the clusters showing statistical significance from paired effect of Wake and N1 on corresponding latency projections. Two clusters were well identified within CP and PTR, respectively. Shifts of latencies were significant in these clusters. Specifically, CP was earlier in N1, as compared to Wake, whereas PTR showed opposite observation.

**Figure 2.**
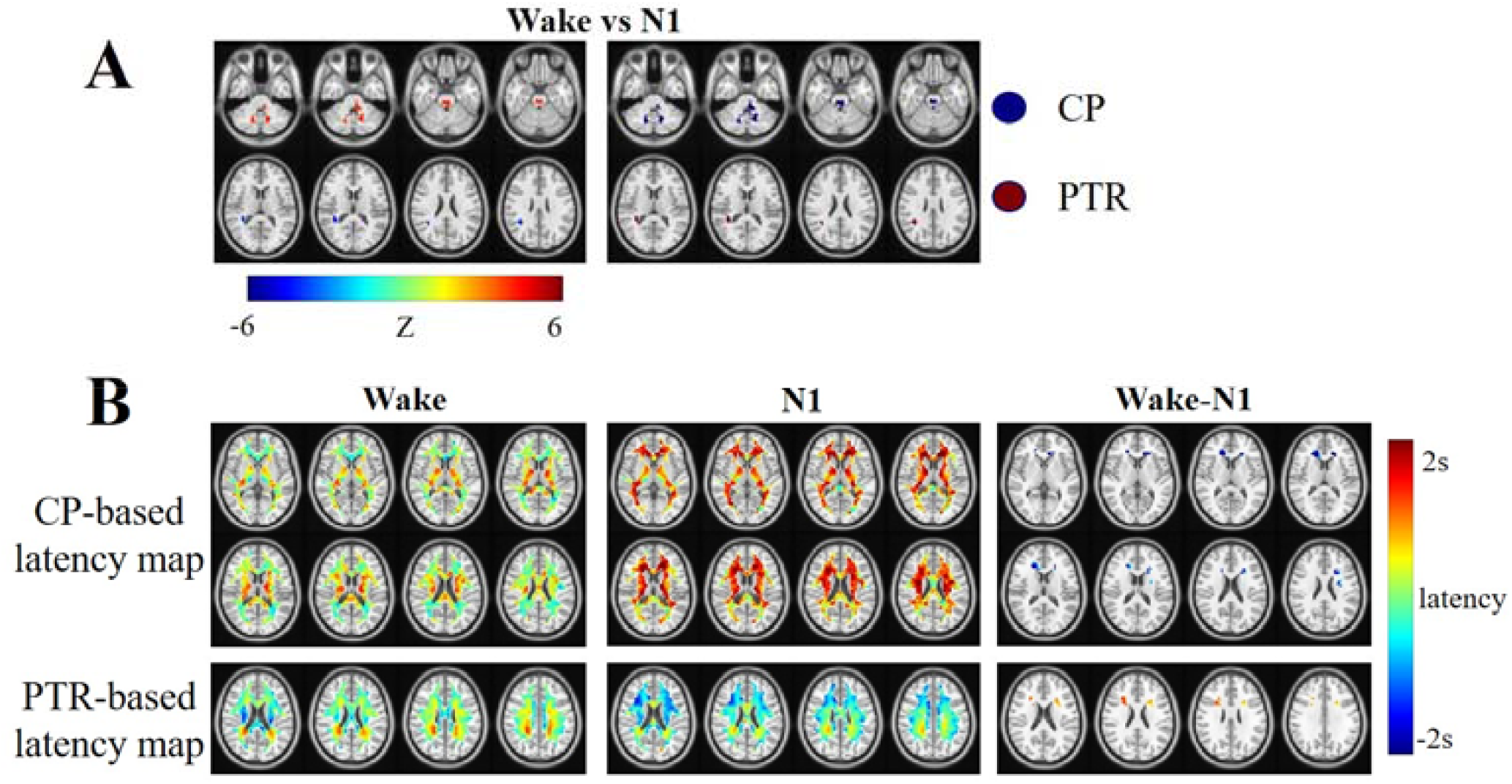
Results of Wake-N1 pair. (A) Left panel: Distributions of clusters, contained within the mask of main effect, showing statistical significance in the post hoc t-test, wherein |Z|>3.29, and p<0.01 with cluster-wise correction. Middle panel: Seeds derived from corresponding clusters in the left panel. Right panel: Labels for corresponding seeds in the middle panel. (B) Left column: Seed-based latency maps of Wake. Middle column: Seed-based latency maps of N1. Right column: Difference seed-based latency maps of Wake and N1 (Wake minus N1). Only clusters showing statistical significance (|Z|>3.29, p<0.01 with cluster-wise correction) were displayed.

Seed-based latency maps (Figure 2-B), wherein value of each voxel represents its latency with respect to the seed, were derived using each cluster defined in Figure 2-A as seed. For CP-based latency map, during Wake, anterior and posterior CR were both found to lead CP, while IC was found to lag CP. This phenomenon drastically changed during N1 in that the remaining parts of WM lagged CP, suggesting a unidirectional path of signal propagation from the inferior to the superior. In PTR-based latency maps, anterior CR (ACR) shifted toward earlier timing (see difference PTR-based latency map in Figure 2-B) when transitioned from Wake to N1.

#### Paired effect of Wake and N2

Figure 3-A displayed the clusters showing statistical significance from paired effect of Wake and N2 on corresponding latency projections. One of the identified two clusters was contained within CP. And the other cluster consisted of retrolenticular part of internal capsule (RPIC) and PTR. When the brain transitioned from Wake to N2, regions in CC and posterior CR (PCR) shifted toward late role. Similar to the observation in paired effect of Wake and N1, the path of signal propagation from the inferior to the superior was also identifiable in CP-based lag map of N2. Besides, major differences between Wake-N1 pair and Wake-N2 pair were the inclusion of voxels from RPIC. In RPICPTR-based latency map, both ACR and posterior CR were found to shift toward earlier timing (see difference RPICPTR-based latency map in Figure 3-B).

**Figure 3.**
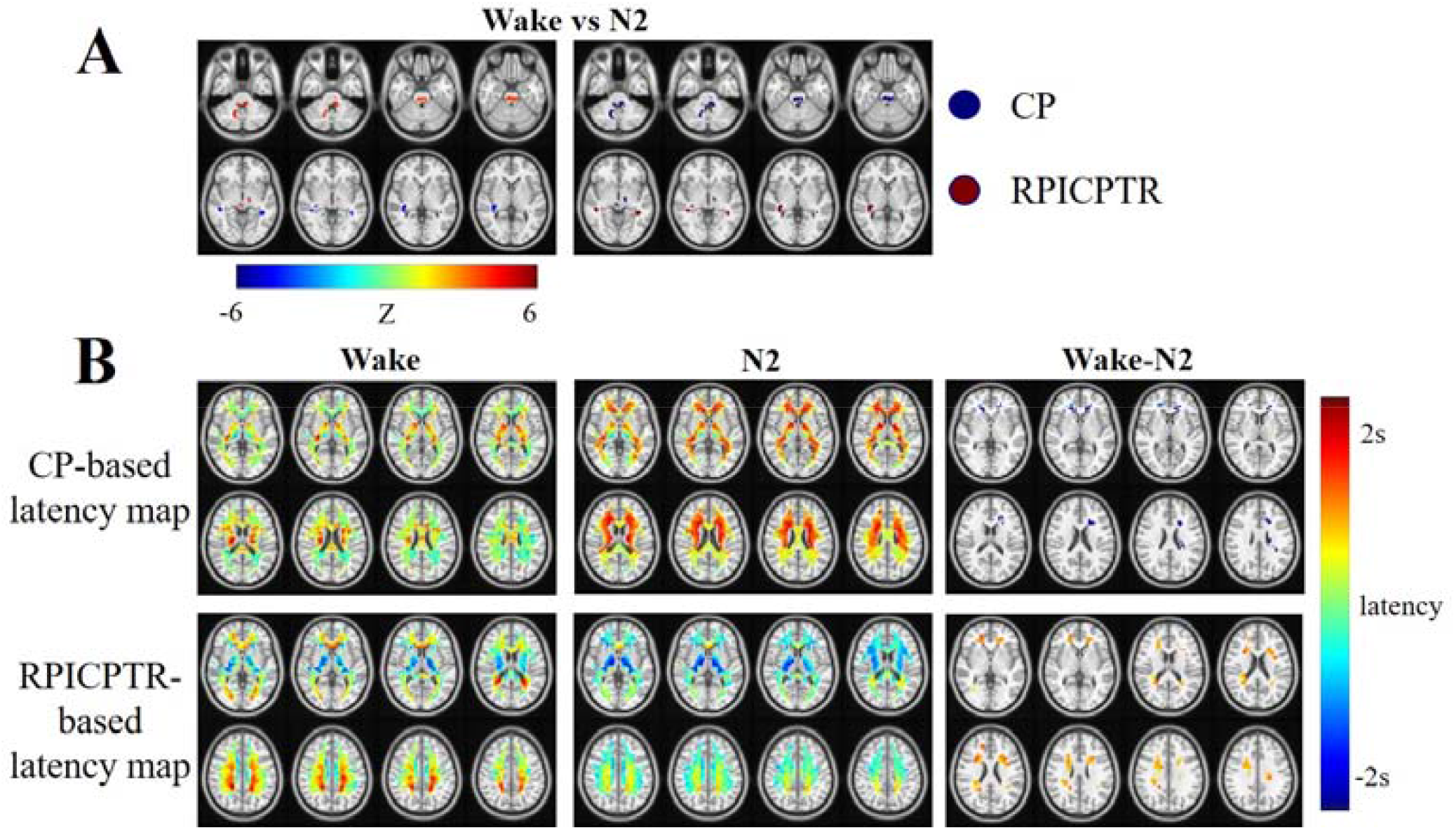
Results of Wake-N2 pair. (A) Left panel: Distributions of clusters, contained within the mask of main effect, showing statistical significance in the post hoc t-test, wherein |Z|>3.29, and p<0.01 with cluster-wise correction. Middle panel: Seeds derived from corresponding clusters in the left panel. Right panel: Labels for corresponding seeds in the middle panel. (B) Left column: Seed-based latency maps of Wake. Middle column: Seed-based latency maps of N2. Right column: Difference seed-based latency maps of Wake and N2 (Wake minus N2). Only clusters showing statistical significance (|Z|>3.29, p<0.01 with cluster-wise correction) were displayed.

#### Paired effect of Wake and N3

Figure 4-A displayed the clusters showing statistical significance from paired effect of Wake and N3 on corresponding latency projections. One of the two identified clusters was contained within RPIC, which shift toward late when transitioned from Wake to N3. The other cluster located in PCR, wherein voxels shifted along the reversal direction against that of the first cluster. Regions in ACR, PTR, and PCR showed shortened latencies w.r.t. RPIC when transitioned from Wake to N3. However, regions in ACR and PTR showed enlarged latencies w.r.t. PCR (difference PCR-based lag maps of Wake and N3 in Figure 4-B).

**Figure 4.**
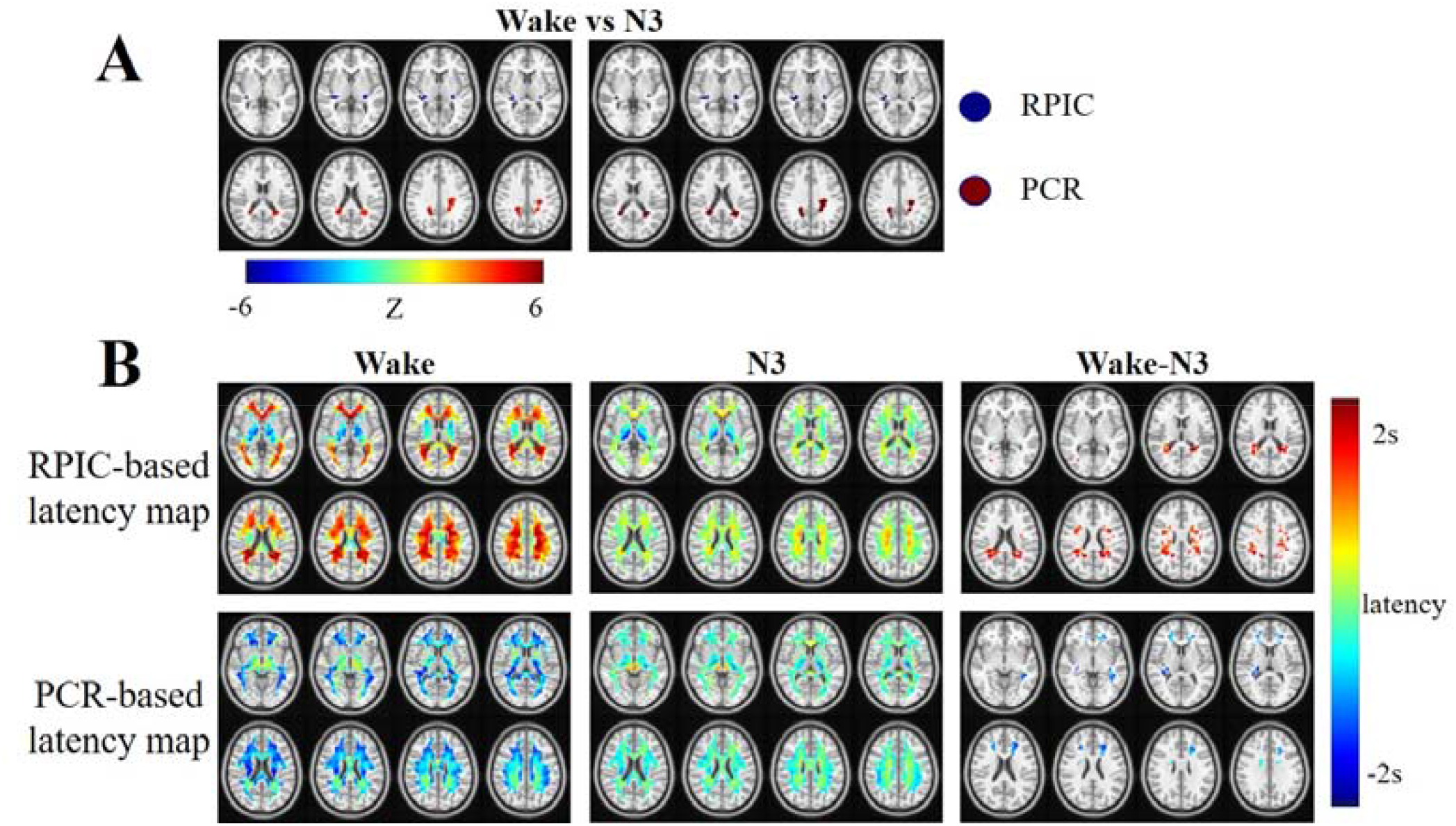
Results of Wake-N3 pair. (A) Left panel: Distributions of clusters, contained within the mask of main effect, showing statistical significance in the post hoc t-test, wherein |Z|>3.29, and p<0.01 with cluster-wise correction. Middle panel: Seeds derived from corresponding clusters in the left panel. Right panel: Labels for corresponding seeds in the middle panel. (B) Left column: Seed-based latency maps of Wake. Middle column: Seed-based latency maps of N3. Right column: Difference seed-based latency maps of Wake and N3 (Wake minus N3). Only clusters showing statistical significance (|Z|>3.29, p<0.01 with cluster-wise correction) were displayed.

#### Paired effect of N1 and N2

Latency projections of N1 and N2 were found to be quite similar, with only a cluster within the white matter of cerebellum (WMCB) was found to exhibit statistical significance in the post hoc t-test of the N1-N2 pair (in Figure 5-A). Part of this cluster was also included when contrasting Wake and N1. Shifting toward later timing was found in N2 than in N1 (see left panel of Figure 5-A), resulting in smaller latency values of the rest voxels with respect to this cluster in the CP-based latency map of N2 (see left panel of Figure 5-B). Seed-based latency maps of both states also showed patterns of inferior-to-superior path of signal propagations.

**Figure 5.**
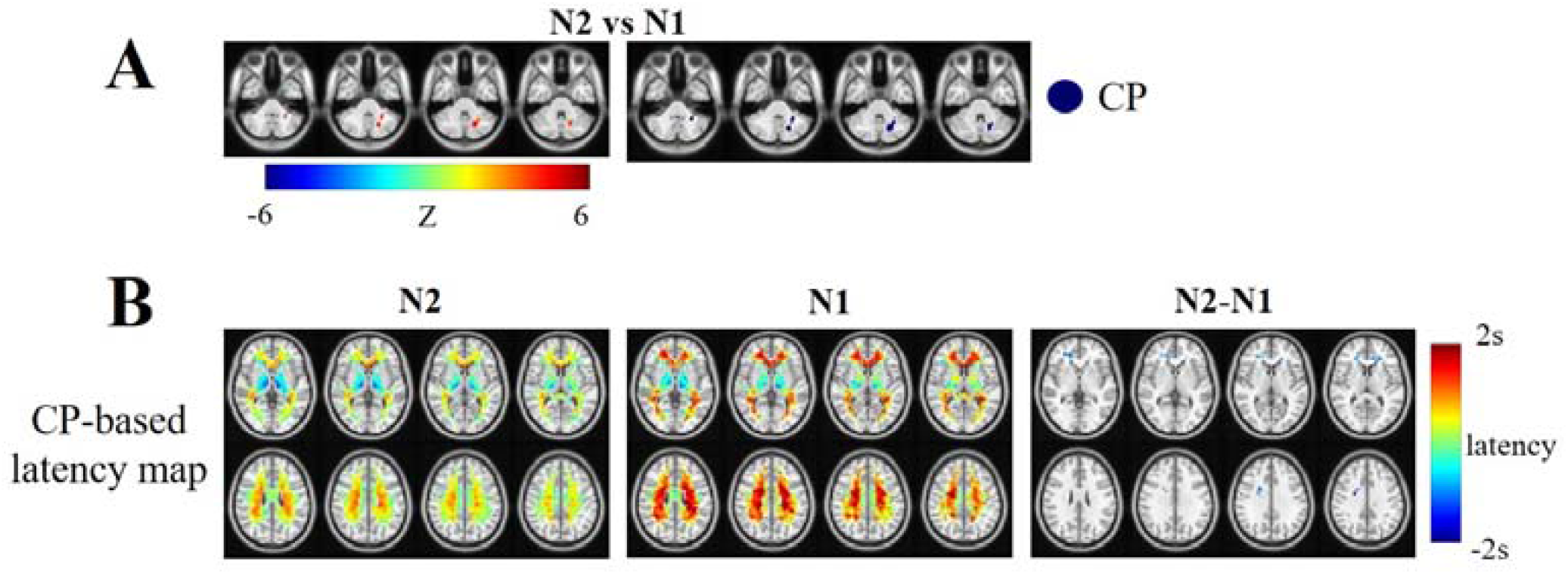
Results of N2-N1 pair. (A) Left panel: Distributions of the cluster, contained within the mask of main effect, showing statistical significance in the post hoc t-test, wherein |Z|>3.29, and p<0.01 with cluster-wise correction. Middle panel: Seed derived from corresponding cluster in the left panel. Right panel: Label for corresponding seed in the middle panel. (B) Left column: CP-based latency map of N2. Middle column: CP-based latency maps of N1. Right column: Difference CP-based latency map of N2 and N1 (N2 minus N1). Only clusters showing statistical significance (|Z|>3.29, p<0.01 with cluster-wise correction) were displayed.

#### Paired effect of N1 and N3

Figure 7-A showed the reorganized clusters of statistical significance within WM regions across N3 and N1 stages. Smaller size of effected regions were identified in the seed-based latency maps of N3-N1 pair than that of N3-N2 pair, as displayed in Figure 7-B. Similar directions of temporal shifting to that in N3-N2 pair were observed in these effected regions. However, regions in GCC showed no significant difference between N3 and N1, which is different from the result of N3-N2 pair.

**Figure 6.**
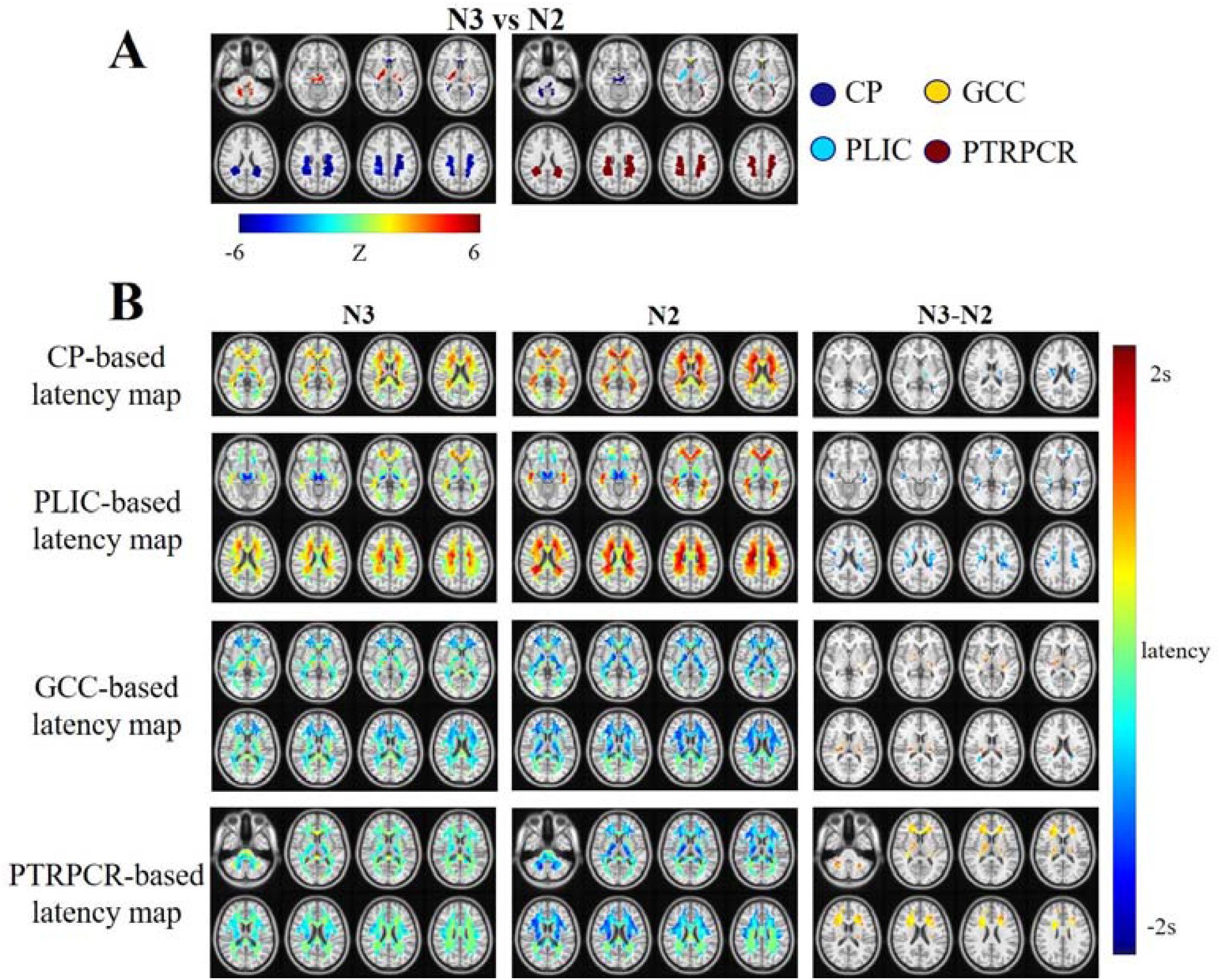
Results of N3-N2 pair. (A) Left panel: Distributions of the cluster, contained within the mask of main effect, showing statistical significance in the post hoc t-test, wherein |Z|>3.29, and p<0.01 with cluster-wise correction. Middle panel: Seeds derived from corresponding cluster in the left panel. Right panel: Label for corresponding seed in the middle panel. (B) Left column: Seed-based latency maps of N3. Middle column: Seed-based latency maps of N2. Right column: Difference seed-based latency maps of N3 and N2 (N3 minus N2). Only clusters showing statistical significance (|Z|>3.29, p<0.01 with cluster-wise correction) were displayed.

**Figure 7.**
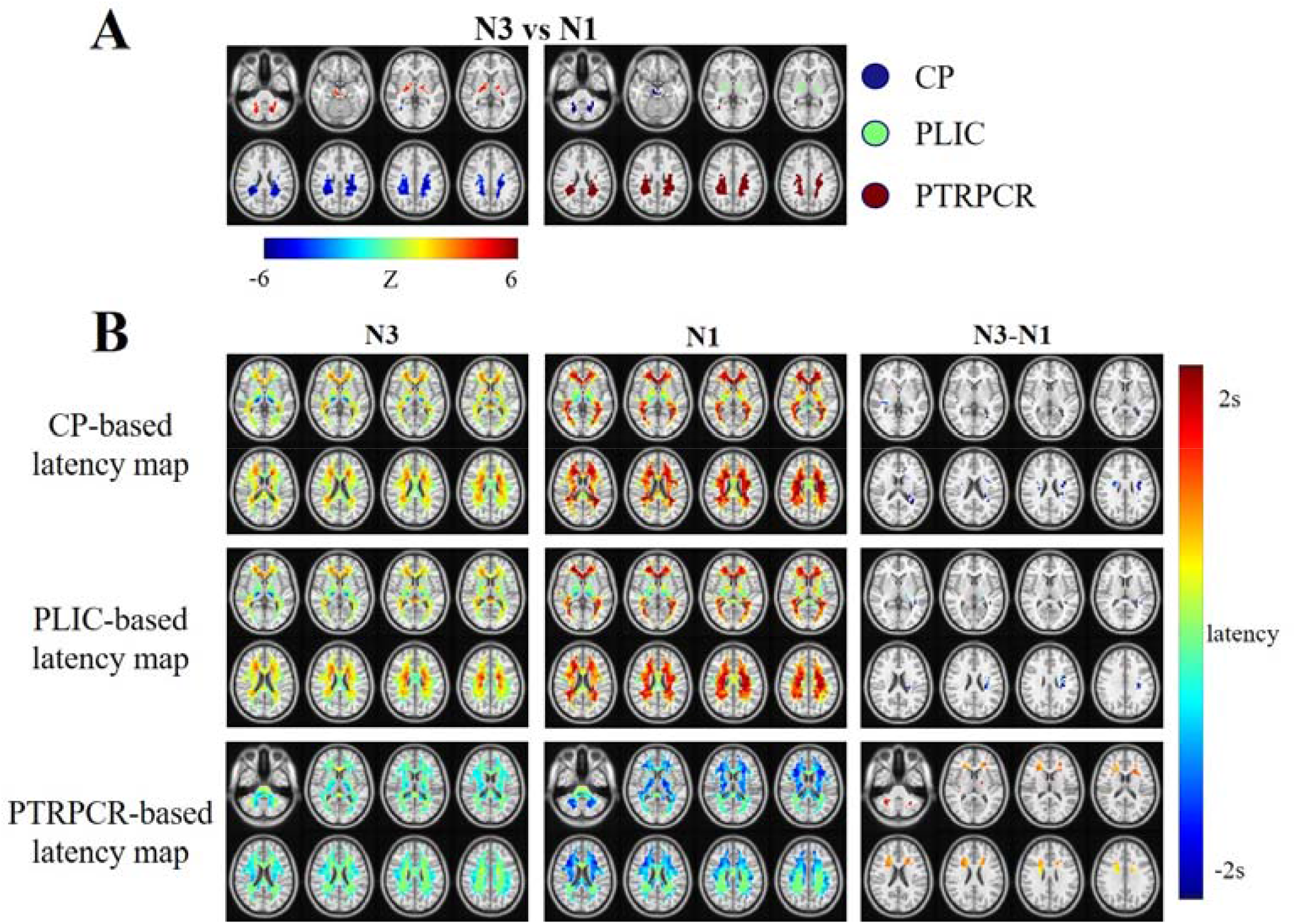
Results of N3-N1 pair. (A) Left panel: Distributions of the clusters, contained within the mask of main effect, showing statistical significance in the post hoc t-test, wherein |Z|>3.29, and p<0.01 with cluster-wise correction. Middle panel: Seeds derived from corresponding clusters in the left panel. Right panel: Label for corresponding seeds in the middle panel. (B) Left column: Seed-based latency maps of N3. Middle column: Seed-based latency maps of N1. Right column: Difference seed-based latency maps of N3 and N1 (N3 minus N1). Only clusters showing statistical significance (|Z|>3.29, p<0.01 with cluster-wise correction) were displayed.

#### Paired effect of N2 and N3

Figure 6-A showed the reorganized clusters of statistical significance within WM regions across N3 and N2 stages. Four clusters were well identified within CP, PLIC, GCC, PTR and PCR (PTR and PCR will be combined as one seed named PTRPCR in later analysis), respectively. Early clusters (CP, SS, and IC) in N2 shifted toward late in N3 while late clusters (GCC and PTRPCR) in N2 shifted toward early in N3, thus yielding a shorter temporal range in N3 than in N2, suggesting faster speed of signal propagations within WM in N3 stage. Despite of this, two more observations were of interests. First, in descent to N3 stage, the above mentioned (see CP-based latency map in Figure 2-B) inferior-to-superior path of signal propagation can be identified from the CP-based latency maps of both N2 and N3. Second, a narrower temporal support of the latency structure of N3 than N2 (Figure 1-A) did not ensure early shift in each of the seed-based latency maps in N3. Exceptionally, in the GCC-based latency map in Figure 6-B, both early (regions in PTR) and late (regions in IC) latency shifts were observed, suggesting inconsistent and stage-specific shifting of latencies.

To summarize, it was evident that the variations of latency projections across wakefulness and NREM sleep stages were underlain by temporal reorganizations manifested as not consistent but various shifts of latencies including both early and late with respect to paired stages, among voxels, which occurred in both WM and GM (Mitra et al. 2015).

### Latency among RSNs in relation to different sleep stages

K-means, a clustering technique, was applied on voxel-wise FC matrix within WM to obtain RSNs. A path of signal propagation, dominated by the direction from the inferior to the superior, was identified by performing latency analysis on WM RSNs, similar to previous study on a different dataset with large cohort of youngsters (Guo et al. 2021a) at the resolution of 8 RSNs. Hence, in this study, the whole WM were parcellated into 8 clusters, as shown in Figure 8-A, by applying K-means on group-level FC matrix consisting of paired voxels of N2 group where the most subjects were included.

**Figure 8.**
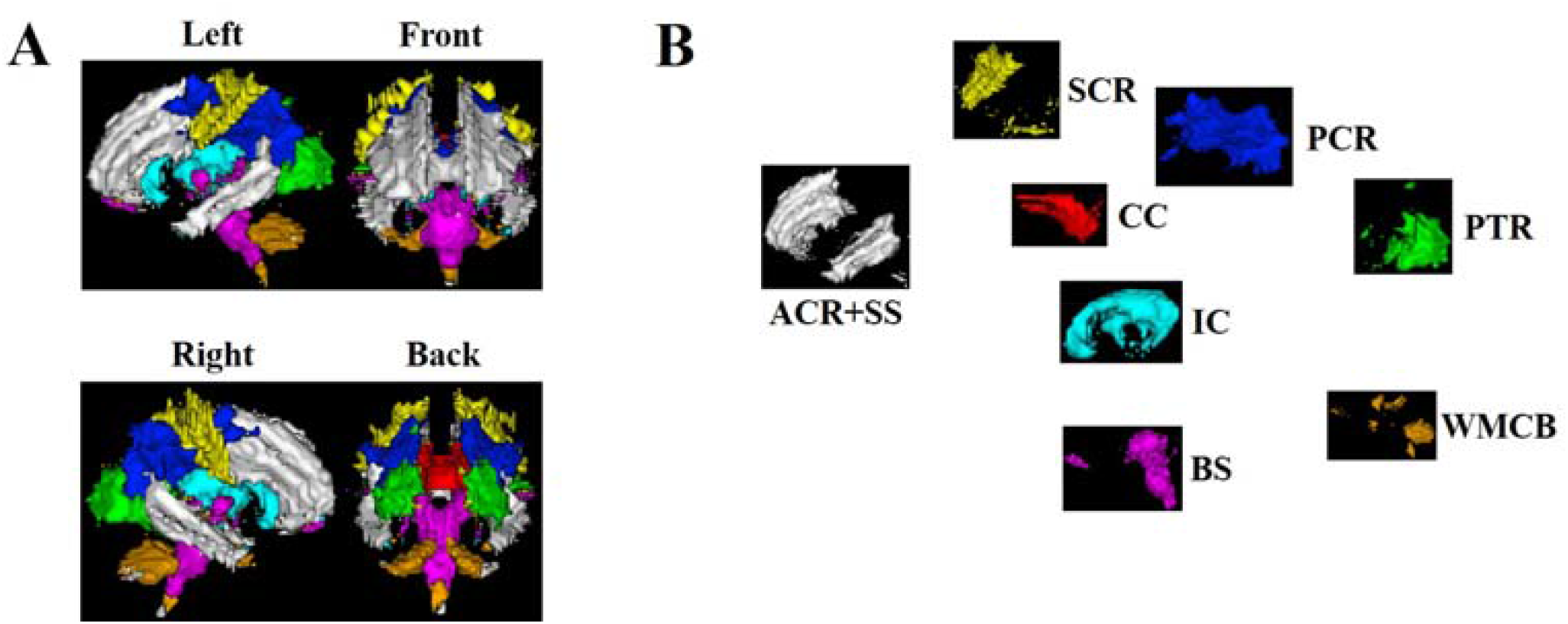
(A) Eight functional networks clustered from paired voxels of group level FC matrix from N2 group. (B) Eight separated RSNs, allocated according to their relative spatial position.

Latency analyses were performed on the RSNs, including corpus callosum (CC), PTR, WMCB, superior corona radiata (SCR), IC, brainstem (BS), PCR, anterior corona radiata and sagittal striatum (ACR+SS). Each RSN was named according to the largest one or two fiber bundles it contains (Figure 8-B).

#### Inter-RSN temporal organizations across wakefulness and NREM sleep stages

Figure 9-A showed the intra- and inter-RSN latencies of paired voxels across Wake and NREM sleep stages. Voxels were ordered according to their corresponding stage-specific latency structure.

**Figure 9.**
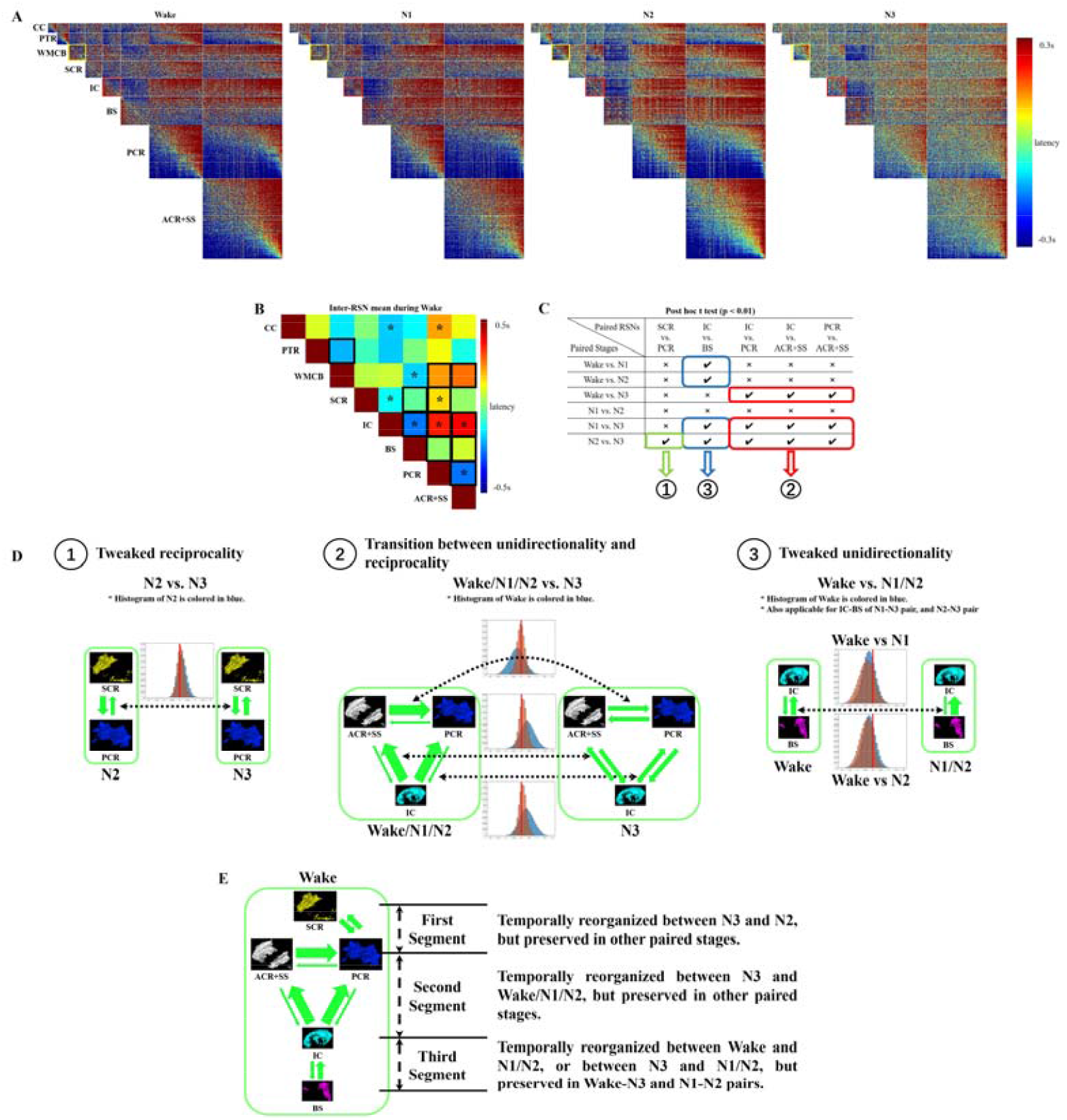
(A) Intra- and inter- RSN latency matrices of WM across different sleep stages. Blocks in yellow and red are referred to in the section of Discussion. (B) Mean inter-RSN latencies during wakefulness. Blocks with asterisks indicated significant (p<0.01, Bonferroni corrected by N=28, i.e., number of the triangular latency blocks in (A)) non-zero values. Blocks in black indicated mean inter-RSN latencies with significant (p<0.01, Bonferroni corrected by N=28, i.e., number of the triangular latency blocks in (A)) main effect of stages. Blocks in black rectangles, along with asterisks in them, were selected for further analysis of paired effect of stages, to investigate the actual paired stages that yielded the inter-RSN temporal reorganizations. (C) Results of the post hoc t-test analysis on the selected blocks. Three types of inter-RSN temporal reorganizations can be identified from the statistical results. (D) Three types of temporal reorganizations between RSNs of (C) in WM. Red vertical line in the histograms indicated value of zero. (E) Temporal organizations among the RSNs in (C) during wakefulness. Different segments of the propagation path reorganized temporally according to the stage pair between which the brain transitioned.

In Figure 9-A, off-diagonal blocks represented the inter-RSN temporal organizations. Some of these blocks showed significant non-zero mean inter-RSN latencies (one sample t test, p < 0.01 with Bonferonni correction of N=28) during wakefulness, which were indicated with asterisks in Figure 9-B. Similar to previous study on WM using completely independent dataset (Guo et al. 2021a), results from this dataset also exhibited network-level lead/lag relationships, suggesting existence of dominant direction of signal propagations. We further applied linear mixed-effect analysis on the mean latencies of the off-diagonal blocks to investigate the main effect of stages on the variations of each mean latency. Blocks showing significant variations of mean latencies across sleep stages were contained within black rectangles, as displayed in Figure 9-B. Blocks showing significant non-zero mean latencies, which also significantly fluctuate across stages, were included for post hoc t-test to explore the inter-RSN temporal reorganizations across specific paired stages, thus yielding five eligible blocks, including SCR vs. PCR, IC vs. BS, IC vs. PCR, IC vs. ACR+SS, and PCR vs. ACR+SS. As displayed in Figure 9-C, significant results of the post hoc t-tests (p < 0.01 with no correction since the blocks were selected from significant results of main effect of stages) showed generally different variations of inter-RSN mean latencies between stage pairs, within which three types of inter-RSN temporal reorganizations were identifiable (see Figure 9-D).

The first type (in Figure 9-D-1) is tweaked reciprocality, observed in the block relating SCR and PCR in paired stage of N2-N3. Histogram of the latency values within this block showed obvious reciprocal communications between SCR and PCR in N2. The latency mean within this block tweaked significantly in N3 but with preserved reciprocal communications.

The second type (in Figure 9-D-2) of temporal reorganization was manifested as transition between unidirectionality and reciprocality, as observable among RSNs including IC, PCR, and ACR+SS in paired stages with N3 involved. In Wake/N1/N2, histograms of latency blocks relating either two of these three RSNs exhibited dominant direction of inter-RSN signal propagations. While in N3 the same blocks as above mentioned showed type of reciprocal communications.

The third type (Figure 9-D-3) of temporal reorganization, named as tweaked unidirectionality, was observed in the latency block relating IC and BS. Histogram of this block during Wake showed a dominant direction of signal propagation from BS to IC (i.e. the inferior to the superior). When the brain transitioned from Wake to N1/N2, the dominated direction of signal propagations was enhanced. When the brain transitioned from N1/N2 to Wake, this direction dominance was weakened. In either situation, the dominant direction did not change, thus tweaked unidirectionality when the brain transitioned between effected paired stages.

To summarize, During wakefulness, temporal organizations among RSNs, including BS, IC, PCR, ACR+SS, and SCR, were manifested as signal propagations along dominant direction (except for SCR and PCR). However, according to the stage pair between which the brain transitioned, temporal reorganizations within different segments (see Figure 9-E) were affected.

#### Intra-RSN temporal organizations across wakefulness and NREM sleep stages

Intra-RSN temporal organizations were shown in the diagonal blocks of latency matrices in Figure 8-A. During wakefulness, RSNs including CC, PTR, SCR, PCR, and ACR+SS, exhibited well-ordered temporal organizations, consisting of early, middle, and late components. Also, the intra-RSN temporal roles were observed to follow that derived from the latency projection. However, reversal latency patterns against that derived from the latency projection, as displayed in WMCB, and IC (blocks in yellow, and red, respectively in Figure 9-A), were also identifiable. Similar reversed intra-RSN latency patterns were also observed in previous study using different dataset (Guo et al. 2021). These reversed latency patterns suggested that temporal orders, derived from the latency projection, of the voxels within the RSN were at least partially reversed against the temporal orders derived from projection of each RSN. However, starting from N1, these two RSNs, showing no reversed latency patterns, progressively exhibited tendency of well-organized temporal orders. Of all the RSNs, BS was found to exhibit the most complicate intra-RSN temporal organizations across the sleep stages in that during Wake, reversal intra-RSN temporal organizations against that derived from the latency projection was identifiable. However, upon the descent of the brain into deeper sleep stages, the intra-RSN temporal orders within BS showed no easily recognizable organizations. More experiments are warranted for further investigations of these observations.

## Discussion

In this work, we applied cross-covariance based latency analysis on fMRI dataset, to study the temporal reorganizations within WM across wakefulness and NREM sleep stages. Latency analysis at the projection level found that different regions in WM exhibited stage-specific reorganizations of temporal orders. Using the regions showing statistical significance in contrasted paired stages as seeds, we further performed seed-based latency analysis to evaluate the relative temporal alterations between each seed and remaining voxels within WM. Seed-based latency analysis suggested that regions in WM shared inconsistent shifting directions of latencies across stages in that some regions were observed to shift toward early while others late. Latency analysis on RSNs, acquired by applying K-means clustering technique on voxel-wise FC matrix, was also performed to investigate intra- and inter-RSN communications across different stages. In general, RSNs in WM exhibited similar reciprocal communications to that in GM, despite of which a dominant direction from brainstem located in inferior WM, to IC located in middle WM, and further up to CR (including ACR+SS, and PCR) located in superior WM, was identifiable in Wake. Also, depending on the stage pair between which the brain transitioned, three types of temporal reorganizations among these RSNs were recognized, i.e., tweaked reciprocality, tweaked unidirectionality, and transition between unidirectionality and reciprocality.

### Physiological sources of the latency structure within white matter

Previous study (Guo et al. 2021a) based on a large cohort of youngsters has demonstrated the existence of highly reproducible latency structure in WM, variations of which were associated with different visual states (eye closed vs eye open, and also eye closed vs. eye open with fixation). However, concerns linger about the physiological sources that underlie the latency structure in WM. A most mentioned potential source relates with a recent finding of sympathetic responses in giving rise to large changes of BOLD signals in WM (and in GM) with a significant time delay (Özbay et al., 2018). The delay is well consistent with the perfusion time between brain regions, and thus the pial artery contractions caused by the extrinsic sympathetic innervation (Lincoln et al. 1995; Hamel et al. 2006) has been modeled as the source driving such fMRI changes (Özbay et al., 2018), which was presumed to further yield latency structure.

It should be noted that progression of NREM sleep into deep stages was accompanied by shifting of automatic balance from sympathetic to parasympathetic dominance (Trinder et al. 2001), based on which non-specific regions showing significant variations of latencies between paired stages are expected to be activated due to changes of systematic sympathetic tone over upper stream extracranial arteries. It is hard to imagine results of different activated seeds among latency projections from different paired stages, e.g. GCC was activated in the paired stage of N3 and N2 (Figure 6-A) but silent in N3 and N1 pair (Figure 7-A). More difficult to conceive were the systematic change of sympathetic tone in producing different directions of latency shifts in the difference seed-based latency maps, e.g. CP and PTR in the right column of Figure 2-A.

Therefore, we further argue that sympathetic tones, if any, were manifested as dynamically modulated surface covering the entire surface of cerebral cortex (and also the surface of cerebellum), wherein the tones of each position were modulated by downstream underlying neural activities from WM (and GM). Actually, several studies have also observed metabolic phenomena in WM, suggesting blood flow from the upper stream pial artery has physiological relevance to functional activity in WM. Early in the century, distributions of oxygen extraction fraction were found to be quite uniform across the brain parenchyma (Raichle et al. 2001), confirming the existence of similar metabolic activity in WM to GM. Recently, the baseline of glucose level in WM in addition to GM was observed to be higher in an EO than in an EC state (Thompson et al. 2016), suggesting the association between variations of aerobic metabolisms in WM with different physiological states. A more direct link between neural activity in WM and local metabolisms was evidenced by the correlated resting state functional connectivity and glucose uptake rate found in WM (Guo et al. 2021b), which establishes a relationship between the functional activity of WM and its metabolic demand. Therefore, it is likely that variations of sympathetic tone in the pial arteries provide a regulatory mechanism to meet varying metabolic demands from deep WM.

### Reorganized latency structure in relation to EEG findings

To better understand the implications of our results, we need to incorporate the findings in EEG wherein the regulation of sleep-wake cycle were mostly investigated (Picchioni et al. 2013; Brudzynski 2021).

From our latency analysis at the RSN level, a primarily ascending path of signal propagation was well identified (Figure 9-E). This path can be divided into two segments, one between BS and IC, and the other between IC and regions in CR (ACR+SS, SCR, and PCR consisted the majority of CR).

In the first segment, BOLD signals were found to propagate from BS, covering the major part of brainstem, to IC, covering the major part of internal capsules. In previous EEG studies, two systems, the brainstem reticular formation (BRF) and thalamus, are indispensable in modulatory control of brain activity across wakefulness and sleep.

BRF is a diffuse and extensive projection systems ascending from the brainstem and directly or indirectly changing ongoing activities in the entire brain via projection fibers (Brudzynski 2021). It plays a critically important role in maintaining and stimulating of wakefulness since large lesions of the BRF yielded coma in animals and humans (Lindsley et al. 1950; Plum et al. 1980; Parvizi et al. 2003). Stimulations of BRF were shown to trigger waking electrical activity in the cortex of sleeping or drowsy animals, resulting in the changes of cortical EEG signals to go from the high-voltage, slow (delta wave, <4Hz), and regular activity of deep sleep, to the low-voltage, fast (gamma, >30Hz), and irregular activity of waking (Moruzzi et al. 1947). Two directions of neuronal projections exist in BRF, descending reticulo-spinal projections facilitating movement and postural muscle tone with multiple behaviors during wakefulness (Peterson et al. 1979; Siegel et al. 1979) and ascending projections into the forebrain which stimulated cortical activation (Steriade et al. 1982; Jones et al. 1985; Jones 1995). On the other hand, during wakefulness, the “resting” brain is not a quiescent but a mainly intrinsic model actively in collecting and interpreting information from the environment and in preparing potential responses to it (Raichle 2010, 2011). This principle also holds true when the brain descends to sleep stages (Seth et al. 2005). Therefore, subjects in resting state fMRI, regardless of wakefulness or sleep, rest quietly with seldom moving or changing the muscle tone. Hence, the primary neuronal projections propagate along the ascending path during resting state fMRI. Importantly, in our latency analysis of the first segment of BOLD signal flow, propagation path from brainstem superiorly to internal capsule, coincided with that identified in above said EEG findings. Also this propagation path persisted across wakefulness and sleep stages by exhibiting tweaked unidirectionality, which seemingly correspond to the results revealed by EEG studies mentioned above.

The second segment related with IC and CR. Signal propagation in this segment was primarily unidirectional, from IC superiorly to CR, during wakefulness, which is well preserved across Wake, N1, and N2 stages. But upon descent to N3, it changed to exhibit reciprocal communication between IC and CR, and also between ACR+SS and PCR, suggesting different working mode of WM RSNs in N3. To relate with the findings from electrophysiological studies, in N3 stage, thalamocortical neurons were found to be hyperpolarized (Franks and Lieb 1994; Franks 2008). Hence, compared with that in Wake, less environmental stimuli were able to reach the cortex via thalamocortical projection fibers contained within corona radiata. On the other hand, electrophysiological signals were found to reciprocally communicated between cortex and subcortical regions, e.g. for memory consolidation in N3 (Sirota et al. 2008; Buzsáki et al. 2013), only to enhance the signal propagations from CR to IC, which may explain the phenomenon that signal propagations between CR and IC changed from primary unidirectionality to reciprocality between stage pair of Wake/N1/N2 and N3.

As discussed above, temporal organizations of WM BOLD signals appeared to relate with electrophysiological signals across wakefulness and sleep stages. Due to few studies exploring the relationship between BOLD and electrophysiological signals from WM, a brief discussion of the relevance between BOLD and electrophysiological signals in GM may help pinpoint future studies in WM. In GM, the independent components of the infra-slow EEG (0.01–0.1 Hz) and of BOLD fluctuations are correlated, pointing to a potentially direct relation between the BOLD and EEG signal at rest (Hiltunen et al. 2014). Also, for latency analysis of GM, hippocampus-based latency maps based respectively on BOLD signals and infra-slow activity from electrocorticography (ECoG) were found to share similar latency structure at the RSN level (Mitra et al. 2016), suggesting a similar spatiotemporal organizations of the two modalities of signals at the RSN level. Providing a more direct evidence (Mitra et al. 2018), experiments based on concurrent imaging of calcium and absorbance of hemoglobin on mice revealed highly similar state-dependent apparent propagation within the whole-cortex with correspondence only limited to infra-slow (<0.1Hz) brain activity, wherein direction of the apparent propagations between visual and motor cortex was further found to coincide with that derived from the infra-slow signals of EEG. These findings suggested infra-slow brain activity to be neurophysiological process which is also reflected in fMRI blood oxygen signals. But at the same time, we also need to point out, in spite of the relevance between GM BOLD and electrophysiological signals, BOLD signals from WM and GM shared different sources. For WM, a possible cell type may be astrocyte, which abounds in WM and was found to functionally relate with health and disease (Lundgaard et al. 2014), and also sleep (Poskanzer et al. 2011). Further studies could focus on animal experiments, exploring the correlations between EEG signals from WM and BOLD fluctuations.

### Clinical implications of WM latency structure

It is widely known that WM is responsible for transducing neuronal information. Intact structure of WM was found to play a critical role in maintaining the inter-hemispheric functional integration of GM (Roland et al. 2017). Also, WM has important implications to studies of learning, cognition and psychiatric disorders (Fields 2008). Structural deteriorations of WM yielded altered FC between WM and GM (Huang et al. 2020), and also between GM and GM (Tahedl et al. 2018; Petsas et al. 2019). Hence, latency analysis may help identify potential biomarkers for WM disease especially that relates with myelination, e.g., multiple sclerosis. Similarly, recent study of Alzheimer’s Disease (AD) showed that degenerations of myelin are potentially associated with age-related deficits in memory (Wang et al. 2020), wherein analyzing the WM latency might add to the repertoire of potential biomarkers for early diagnosis of AD. In fact, previous study (Mitra et al. 2015b) revealed the potential causal role of temporal organizations in giving rise to RSNs, based on which biomarkers relating with latency structure may be more fundamental than conventional FC. Moreover, structural intactness of WM was found to relate with sleep duration, cognitive performance (Grumbach et al. 2020), and further to exert influences on the neuronal synchrony during sleep (Erlan et al. 2019). Future experiments could be extended to explore the altered latency structure of WM, along with its changed temporal role in relation to the latency structure of GM, to reveal the potential alterations of inter-RSN functional integrations upon deteriorations of WM structure in sleep.

## Author Contributions

Bin Guo, Qihong Zou, and Jia-Hong Gao conceived the project and research approach. Bin Guo designed the methods, implemented the methods and performed the data analysis. Bin Guo, Guangyuan Zou, and Jun Jiang processed the data. Bin Guo wrote the paper; Fugen Zhou, Guangyuan Zou, Jun Jiang, Qihong Zou, and Jia-Hong Gao reviewed and edited it.

## Acknowledgements

This work was supported by the National Key Research and Development Program of China (2018YFC2000603 and 2017YFC0108901); the National Natural Science Foundation of China (81871427, 81671765, 81790651,81790650, 81727808, 81430037 and 31421003); the Beijing Municipal Science & Technology Commission (Z181100001518005, Z161100002616006 and Z171100000117012); the Beijing United Imaging Research Institute of Intelligent Imaging Foundation (CRIBJZD202101). We thank the National Center for Protein Sciences at Peking University in Beijing, China, for assistance with MRI data acquisition and data analyses.

